# The Coevolution of Social Learning and Sensitivity to Changing Environments

**DOI:** 10.1101/080507

**Authors:** Richard McElreath

## Abstract

There is widespread agreement that social and individual learning are adaptations to varying environments. However, existing theory assumes that organisms cannot detect changes in the environment and instead adapt to averages. This paper develops the first analytical model that allows for the simultaneous coevolution of socially learned traditions, reliance on social learning, and signal detection for environmental change. There are numerous conditions under which detection can be stable once common but cannot invade the population when rare. When signal detection is maintained by selection, it always leads to pure separating equilibria at which organisms always learn individually when they believe the environment has recently changed and otherwise always learn socially. Detection can increase mean fitness at equilibrium, but it may also reduce it.

## 1. Introduction

This paper asks when natural selection will favor an organism’s paying a cost to detect changes in the environment, provided that cues of environmental change adjust use of individual and social learning. I use formal modeling to address this question. But the motivation for the paper is really empirical, meant to address a gap between the structure of the theory and how it is interpreted in light of data. When I was a new assistant professor, I set out with colleagues Peter J. Richerson, Mark Lubell and several industrious PhD students to follow Kameda and Nakanishi (2002) and develop an experimental program for studying the adaptive design of social learning in humans (McElreath *et al.* 2005, 2008, Efferson *et al.* 2007, 2008). The goal was to evaluate the predictions of theory, by using different experimental treatments to simulate differences in theoretical parameters, such as rate of environmental change, that lead to changing predictions for reliance on social learning. A number of other laboratories have also explored the same sorts of questions, and results confirm the qualitative agreement between what the models predict and how real people (or at least, real university students) change their social learning strategies in response to changes in aspects of the social and physical environment (Morgan *et al.* 2012, provide a recent and clear example).

But a number of issues with interpreting the experimental results have preoccupied us. For example, laboratory social learning experiments rarely create the differences in experience that both natural communities and theoretical models always possess. Instead, a group of naive participants are asked to learn from one another, and their behavior is predicted using models that assume a different life history, in which naive individuals always coexist with experienced individuals. As a result, those rare experiments that do established “overlapping” generations may show much more powerful and realistic social learning effects (Baum *et al.* 2004, Jacobs and Campbell 1961).

Another concern, and the one that occupies the remainder of this paper, has been that the formal theory itself does not allow for the kind of savvy attention to contextual variables that our design and interpretation of the experiments assumes. We have been expecting savvy context-sensitive deployment of individual and social learning strategies, based upon the interpretation of formal models in which learning strategies respond to context only over evolutionary time (reviewed in McElreath *et al.* in press). Even the interpretation of my own fieldwork has used this slight of hand (McElreath 2004). There are a few cases in which models have allowed for contingent strategy use (Henrich and Boyd 1998, Boyd and Richerson 1996, Enquist *et al.* 2007, McElreath *et al.* 2008). However, for the most part the literature has focused on how evolution, rather than individuals, could strategically adjust learning.

This focus has made it difficult to really say what theory predicts. It makes sense to view the evolution of contingent social learning as a special case of the general theory of phenotypic plasticity. Social learning is itself a form a phenotypic plasticity, and the evolution of plastic use of it is a kind of meta-plasticity. We might wonder when such meta-plasticity might evolve, because the general evolutionary ecology literature has long confirmed that phenotypic plasticity is not always favored by natural selection (Levins 1968, DeWitt *et al.* 1998). The general literature on the evolution of phenotypic plasticity is too vast to review here, but it is worth noting that selection may not favor an organism’s adjusting phenotype in response to cues (e.g. Cohen 1967, Getty 1996, Tufto 2000), and non-contingent strategies can be favored even when environmental cues are reliable (McNamara and Dall 2011). These results have been recently generalized to a wide range of biological phenomena (Altenberg 2012, in press). Some of my own previous theory on the evolution of social learning has turned out to illustrate it (McElreath and Strimling 2008, as cited in McNamara and Dall 2011). On the other hand, in a recent high-profile simulation tournament of the evolution of social learning strategies, one of the most successful strategies regulated social learning by using the time since behavior was learned, in combination with an estimate of how quickly payoffs change over time (Rendell *et al.* 2010).

In light of these results, it’s worth wondering when we should expect people and other organisms to pay attention to cues that regulate learning strategy. We might begin by reconsidering formal models of gene-culture co-evolution in the presence of sensitivity to changing environments. When will natural selection favor using cues of spatial or temporal environmental change to regulate mode of learning? Ongoing debates about the adaptiveness of strategies such as conformist transmission, which has long been central to the gene-culture coevolution literature (Boyd and Richerson 1985), may depend upon understanding selection for such sensitivity (McElreath *et al.* in press, Nakahashi *et al.* in press). And as the planet warms and is otherwise rapidly altered by human activity, predicting and understanding species’ responses will partially depend upon our ability to make sense of the design of environmental sensitivity (Sih *et al.* 2011).

The rest of this paper develops a first model that directly addresses the question: *When will natural selection favor attention to cues of temporally changing environments in order to regulate reliance on individual and social learning?* I use a common gene-culture or dual-inheritance modeling framework (Cavalli-Sforza and Feldman 1981, Boyd and Richerson 1985). I add to this structure another heritable component of plasticity that invests in detecting temporal changes in the environment. The organism can use different learning strategies depending upon whether or not it believes the environment has recently changed. Using a signal detection framework, like a number of previous theoretical studies of phenotypic plasticity (e.g. Getty 1996), I show that gene-culture coevolution may lead to substantial investments in detecting change, but that such investment is not always favored. Indeed, the range of conditions that can stabilize sensitivity to changing environments is always larger than the range that will allow it to invade the population. But whenever detection does evolve, it leads to a perfect separating equilibrium at which the organism always learns individually, when it believes the environment has recently changed, and otherwise always learns socially, when it believes the environment has not recently changed. The result is that much more social learning is observed, once detection evolves. Despite the increase in the amount of social learning, the expected population growth rate may nevertheless increase in the presence of detection, due to adaptive allocation of individual learning to time periods in which it is needed most. I close the paper by considering limits of the model, avenues for future work, and the impact of these results on the interpretation of empirical evidence.

## 2. Model Assumptions

For comparability to existing theory, I use a traditional discrete generation, infinite population framework to construct the model (Cavalli-Sforza and Feldman 1981, Boyd and Richerson 1985, Rogers 1988). Many models address a similar core problem. The adaptive challenge for the organism is to acquire optimal behavior in a temporally varying environment. Since the optimal behavior changes over time, always learning socially is never evolutionarily stable. But similarly, since asocial learning is more costly, unless optimal behavior changes very quickly, some social learning is usually favored. The rate of environmental change and the cost of asocial learning govern the evolutionarily stable mix of social and asocial learning. Because of geometric mean fitness effects and bet hedging, natural selection tends to favor mixed learning strategies over pure ones (McElreath *et al.* in press, Perreault *et al.* in press).

In this paper, I use continuous strategy spaces, allowing individual genotypes to code for probabilities of individual and social learning in different contexts. I keep as much as possible about the core model the same as other papers. The traditional framework has several drawbacks, which I explore in the discussion. However, I wish to begin by changing as little as possible about existing theory, in order to understand the consequences of allowing sensitivity to changing environments to regulate social learning. I introduce into the basic model the ability for an organism to detect environmental change and use different probabilities of social learning depending upon its inference. I develop a weak selection approximation to the geometric mean fitness of a mutant, which allows me to define the evolutionary dynamics of detection. The rest of this section defines the model in detail.

### 2.1. Population and life cycle

Suppose a large well-mixed population of semelparous organisms that are capable of both individual and social learning. The environment the organisms inhabit is everywhere the same, but may change from one generation to the next. Let *u* be the chance of the environment changing in any given generation. The current state of the environment prescribes a unique behavior that results in an increase in expected reproduction (“fitness”) *b*. All other behavior results in no change in fitness. When the environment changes, it changes to a new state it has never had before, and all previous behavior is rendered non-optimal.

### 2.2. Heritable strategies

Behavior is always acquired via learning. But learning strategy is a heritable trait that specifies the probability of using individual learning, instead of social learning. Employing individual learning means that the organism pays a fitness cost *bk* (a proportion *k* of the maximum gain *b*) for a chance *s* of learning optimal behavior. Social learning means that an individual pays no up-front learning cost relative to individual learning, but instead copies a random member of the previous generation. While cheaper, social learning may or may not yield currently optimal behavior, and so it may ultimately be more expensive than individual learning, especially just after a change in the environment.

The adaptive challenge the model explores is how individuals regulate their learning strategy, based upon information that the environment has recently changed. Let *p*_*s*_ be the heritable probability of deploying individual learning when an individual believes the environment has been *stable*, since the last generation. Let *p*_*c*_ be the probability of deploying individual learning when the individual believes the environment has *changed* since the last generation.

### 2.3. Signal detection

Individuals acquire beliefs about the state of the environment via investment in detecting signals of recent change. These signals may be anything from changes in perceived efficacy of a technology or technique to appreciation of others’ opinions on whether or not the environment has changed. I comment more on the nature of such signals in the discussion. The crucial limiting assumption in this model will be that it is individual investments that affect belief formation. Let *d* be the probability of correctly detecting a change in the environment. This is an individual heritable character, with population mean *d*^⋆^. Let *f*(*d*) be a function that determines the probability of a false positive, of thinking the environment changed when it did not. The population mean rate of false positives is *f*(*d*^⋆^) = *f*^⋆^.

I leave this function undefined for now. However, there are several limiting assumptions we can make about the shape of this function, before defining it, and these assumptions will be sufficient to prove the invasion and stability criteria for the model. The general shape of the function *f*(*d*) comes from analogy to a Receiver Operating Characteristic (Green and Swets 1966). A Receiver Operating Characteristic (ROC) describes the tradeoff between accuracy and error in a classification task. As the ability of a signal or test to detect true cases rises, so too does the rate of false positives. As a result, optimal detection in real classification tasks almost always means accepting some false-negatives as well as false-positives.

Readers familiar with the signal detection literature will recognize *d* as the sensitivity and *f*(*d*) as one-minus-specificity, the Type II error rate. The exact shape of the tradeoff between detection and false alarms depends upon the details of each case, but the general nature of this tradeoff is nearly universal in signal detection. Figure 1 illustrates the general shape *f*(*d*) must take. First, I restrict *f*(*d*) to continuous, differentiable functions. Second, realistic signal detection problems have a few recurring features. The only reliable way to detect all true cases is to always assume that the event has occurred. This implies that *f*(1) = 1—if the detection rate is 100%, then the false positive rate is also 100%. Likewise, the only way to miss every true case is to assume the event never happens, *f*(0) = 0. Third, I assume that *f*(*d*) ≤ *d*, the rate of false positives is everywhere less than the rate of true detection, unless *d* = 1. Finally, the previous assumptions imply that the rate of change in false positives is everywhere positive or zero, *f′*(*d*) ≥ 0, and that the acceleration of false positives is strictly positive, *f″*(*d*) > 0. It is also necessary that *f′*(0) < 1, as a consequence of assuming *f*(*d*) ≤ *d*.

**Figure 1:**
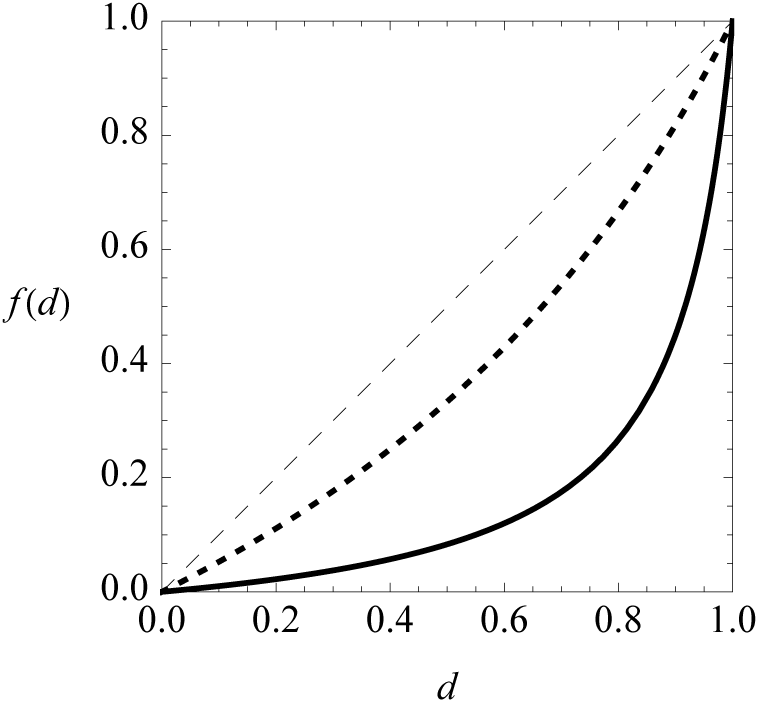
Relationship between true detection rate *d* to false positives *f*(*d*). Increasing investments in *d* on the horizontal axis lead to increases in the rate of false positives, *f*(*d*). The dashed line on the diagonal represents *f*(*d*) = *d*, where the signal is useless because the rate of true positives equals the rate of false positives. Below this line, the signal provides information that allows individuals to detect true changes in the environment. Example curves in this figure are the function *f*(*d*) = *ad/*(1 + *a − d*). The solid curve is for *a* = 0.1. The dashed curve is for *a* = 1. Larger values of *a* indicate higher rates of false positives for any given rate of true positives.

Although I will prove most of the interesting features of this model for any *f*(*d*) that fits the restrictions above, in order to illustrate the dynamics of the model, I will later need a particular function *f*(*d*). Specifically, I will use a flexible hyperbolic function for all numerical examples in this paper:

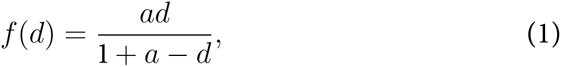

where *a* > 0 is a parameter that determines how quickly *f*(*d*) increases as *d* increases. This function ensures that when *d* = 1 (detection is perfectly accurate), that *f*(1) = 1 also (false positives always occur). At the limit *a* → 0, *f*(*d*) = 0, the signal is perfect. But at the limit *a* → ∞, *f*(*d*) = *d* and the signal is useless, because behaving according to the signal is just like guessing. Figure 1 plots this function for *a* = 0.1 and *a* = 1.

### 2.4. The cost of signal detection

Attempting to detect environmental change carries a fixed fitness cost, *bℓd*, where *ℓ* > 0 is a new parameter that governs the marginal cost of signal detection. Individuals who are increasingly sensitive to environmental change pay an increasing fitness cost. This assumption allows for a wide range of different mechanistic hypotheses. If we suspect that information about environmental change is quite cheap to acquire and process, then *ℓ* can be made to be close to zero. If we suspect instead that such information is costly to acquire or process, then *ℓ* will be large.

### 2.5. Fitness at time *t*

With the assumptions above, we can write a general fitness function for a mutant individual with individual learning probabilities *p*_*s*_ and *p*_*c*_ and detection rate *d* in a population in which everyone else has probabilities 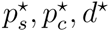. Let *w*_0_ be baseline fitness accrued through other activities. Let *t* be the number of generations since the last change in the environment. For notational simplicity, I define *f* ≡ *f*(*d*) and *f*^⋆^ ≡ *f*(*d*^⋆^). When the environment has changed since the previous generation completed their learning, *t* = 0, detection of true change affects fitness. Then the expected fitness of the mutant is:

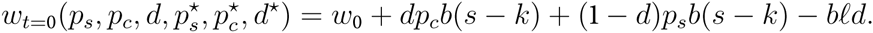

The fitness of this mutant at a time *t* > 0 generations since the last change in the environment is:

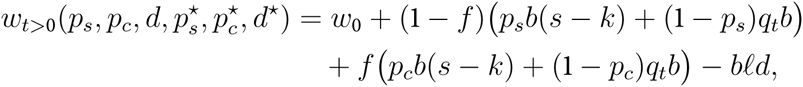

where 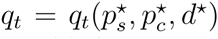 is a function that yields the probability of acquiring optimal behavior via social learning, *t* generations after a change in the environment. I derive *q*_*t*_ in the next section.

All together the fitness of a mutant *t* generations after the most recent change in the environment is given by:

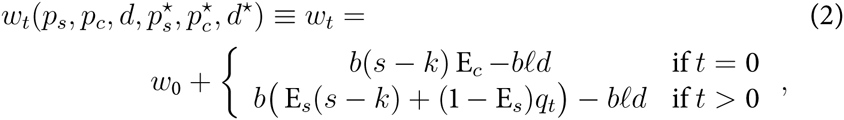

where E_*c*_ = *dp*_*c*_ + (1 − *d*)*p*_*s*_ is a mutant’s expected amount of individual learning, just after a change in the environment, and E_*s*_ = *fp*_*c*_ + (1 − *f*)*p*_*s*_ is the mutant’s expected amount of individual learning when the environment has not recently changed.

### 2.6. Quality of social information, *q*_*t*_

The next step in computing the growth rate of the mutant strategy is to compute *q*_*t*_, the probability of acquiring adaptive behavior via social learning, *t* generations after the most recent change in the environment. The problem is to define the recurrence process by which adaptive behavior accumulates in the population. Just after a change in the environment (*t* = 0), there is no chance of acquiring adaptive behavior via social learning, because all behavior that was learned in previous generations is now non-optimal. Every generation that the environment remains stable, adaptive behavior is pumped into the population via the action of individual learning.

In the Supporting Information, I use the logic above to derive the explicit function for *q*_*t*_, the probability of acquiring adaptive behavior via social learning, *t* generations after a change in the environment:

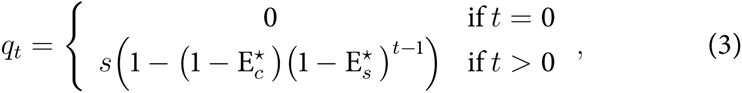

where 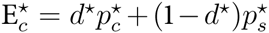 is the average amount of individual learning in the population, just after a change in the environment, and 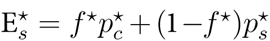 is the average amount of individual learning when the environment has not recently changed. I use this function in the next section to estimate the growth rate of the mutant.

### 2.7. Long run expected growth rate

The probability that the mutant will increase in frequency depends upon the stochastic nature of the environment. To compute the required expression, we note that selection in time varying environments, at least with simple life histories such as these, will maximize the geometric mean fitness, not the arithmetic mean fitness. For a particularly clear explanation of this fact, see Cohen (1966). I label the geometric mean fitness of the mutant 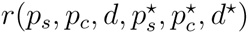 and work with its natural logarithm. This expression is defined as:

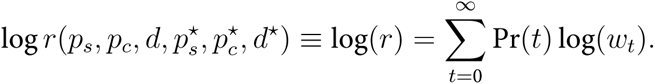

This is just the natural logarithm of the geometric mean fitness of the invading mutant.

The only part of this puzzle still missing is a function defining Pr(*t*), the chance the environmental takes on the state *t* in any given generation. This is given by Pr(*t*) = *u*(1 *− u*)^*t*^. If the environment just changed, then *t* = 0, and this happens with probability *u*, by the definition of *u*. In order to reach *t* = 1, the environment has to remain stable for one generation. The chance of this is *u*(1 − *u*). For *t* = 2, the chance must be *u*(1 *− u*)^2^, because a sequence of two generations without a change is necessary. A similar derivation of this geometric relationship appears in Rogers (1988).

### 2.8. Weak selection approximation

The expression log(*r*) above is inconvenient for analysis. There is no known method for closing this kind of infinite series, in which the index variable *t* is an exponent both inside and outside of the logarithm. To make progress, I use the customary tactic. I construct a weak selection approximation by using a Taylor series expansion of log(*r*) around the point *b* = 0 and keeping the linear term only. This provides an approximation of the model for *b*^2^ ≈ 0, corresponding to the assumption that selection is weak:

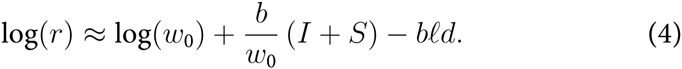

*S* is a term summarizing the fitness benefits of social learning, and *I* is a term summarizing the fitness benefits of individual learning. These terms are:

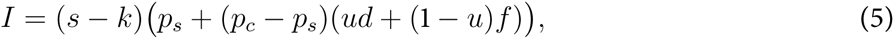

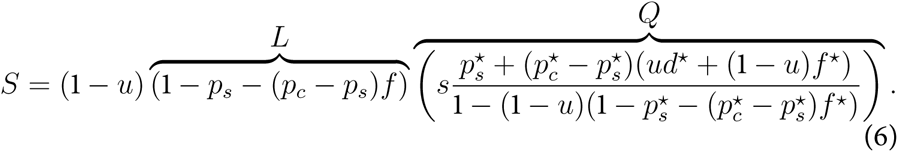

Some sense can be made of these expressions, before analyzing the dynamics. Consider the expression for *I*. It is proportional to *s − k*, the proportion of fitness benefits that remain after subtracting the costs of individual learning. The rest of the expression merely quantifies the mutant’s rate of individual learning, taking into account signals of environmental change and the different rates of learning they create. Note that the common-type trait values 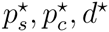 do not appear in the expression for *I*.

Social learning, expression *S*, however does depend upon common-type strategy. Social learning only pays when the environment has been stable for at least one generation, and so the entire expression is proportional to 1 − *u*. The term labeled *L* is the rate of social learning for the mutant, when the environment is stable. The term labeled *Q* quantifies the expected quality of social information. It is exactly the expected probability of acquiring adaptive behavior via social learning, conditioned on the environment being stable (*t* > 0). It depends upon the common type phenotypes 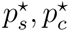 and *d*^⋆^, because the common type creates the cultural environment that the mutant experiences. Note that the numerator of this term is just the common-type rate of individual learning, which is the rate at which new adaptive behavior enters the population. This input of adaptive behavior is discounted by the denominator, which is one minus the probability of social learning, given that the environment is stable. As the amount of social learning increases, the denominator gets smaller, making any inputs from individual learning accumulate more. So the denominator in total can be thought of as a cultural turnover rate. When it is small, because social learning is common and the environment is relatively stable, the entire value of *Q* is increased through accumulation of past innovations. When the denominator is instead small, because either social learning is rare or the environment is relatively unstable, then *Q* is reduced.

## 3. Analysis

To analyze the model, I use a tactic common in evolutionary ecology and evolutionary game theory. If mutants are rare and phenotypically very close to the common type, then the change in each trait is proportional to the rate of change in mutant fitness:

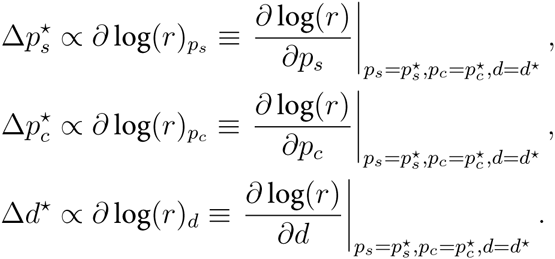

By analyzing these three gradients, it is possible to determine the equilibria and stability conditions of the model.

### 3.1. Equilibria and stability

While this model has no true equilibria, because it is stochastic, it does have steady state expected values for the state variables. It turns out that there are only two possible steady states in this model. Let 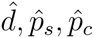 denote expected values of the state variables that satisfy 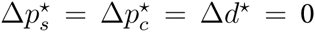. Either the system comes to rest at a detectionless steady state where 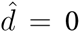 and 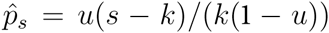, or it comes to rest where 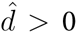 and 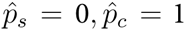. In the Supporting Information, I show how to derive conditions for the stability of both the 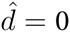 steady state and the 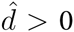 steady state. In the remainder of this section, I present these conditions and try to motivate their logic.

#### 3.1.1. Condition for detection d > 0 to invade

The condition for detection to invade from zero is:

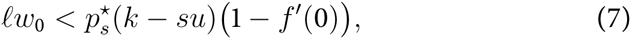

where 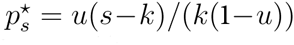 is the probability of individual learning when *d*^⋆^ = 0, and *f′*(0) is *∂f*/*∂d*|_*d*=0_, the initial rate of increase in false alarms of environmental change. This expression confirms the intuition that detection can more easily invade when its direct fitness cost, *ℓ*, is low. Also intuitively, when false positives increase slowly with detection, *f′*(0) is small, detection more easily invades.

#### 3.1.2. Level of detection when 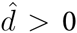

When detection does invade and increase from zero, the learning state variables evolve to 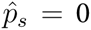 and 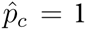. There is no similarly simple expression for the value of 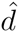. The expression for the steady-state amount of detection is complex, but is defined implicitly by:

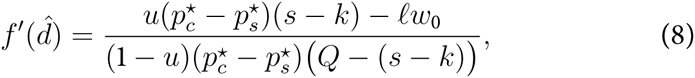

or equivalently, using the fact that 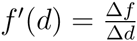:

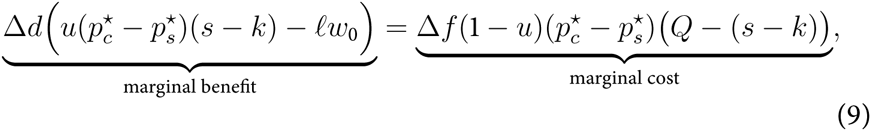

where *Q* has the same form as in Equation 6, quantifying exactly the expected probability of acquiring adaptive behavior via social learning, conditioned on the environment being stable (*t* > 0). Equation 9 states what in hindsight is obvious: selection converges to the value of *d*^⋆^ at which the marginal benefits of detection are equal to the marginal costs of false positives. But it also identifies the precisely relevant marginal benefits and costs, which I believe is less obvious. I’ll unpack this equation one part at a time.

First, notice that everything except the direct cost of detection, *ℓw*_0_, is scaled by the term 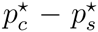. This difference is how much more individual learning is expressed when an individual believes the environment has just changed (*t* = 0). When this difference is zero, detection has no effect on behavior, because learning is not contingent upon the signal. The direct cost *ℓw*_0_ is the unconditional marginal cost of investing in detection. It is unaffected by the difference 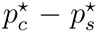, because it is a flat fitness cost that always reduces net benefits.

The left side of Equation 9 summarizes the net marginal benefits of detection. The probability that the environment changes (*u*) is multiplied by the expected net benefit of individual learning, *s − k*. This is the rate of benefit from correct detection. This expected benefit is unconditional on the frequencies of individual and social learning, because when *t* = 0, social learning never pays and individual learning’s fitness is independent of the frequencies 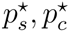. Finally, the marginal cost of detection *ℓw*_0_ is subtracted to yield the net benefit of detection.

The right side of Equation 9 is the marginal cost of detection. This expression quantifies the expected foregone benefits of mistakenly learning individually when the environment is stable (a false positive). *Q* is the probability of acquiring adaptive behavior via social learning (given that *t* > 0), and *s − k* is again the net benefit of individual learning. The difference is the net benefit of social learning, or rather here the net cost of a false positive, which induces an individual to learn individually when it might have learned socially.

#### 3.1.3. Condition for detection d^⋆^ > 0 to be stable

Whether or not Expression 7 is satisfied, it is possible for detection to be stable once common. The condition in this case is a complicated expression that yields little qualitative insight, but I show in the Supporting Information that it can be satisfied even when detection cannot invade.

To begin to understand why this is the case, note that there are three dynamic regimes in the model. Figure 2 illustrates these. Each plot shows the phase plane dynamics of 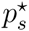 and *d*^⋆^ when 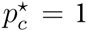. The state variable 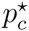 can be fixed at one, because it evolves to one very quickly for most parameter combinations. This allows us to understand the reduced two-dimensional system, as shown in the figure. In each plot, the gray lines with arrows show the flow of the system at each point in the 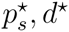 space. The black curve is the 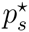 fitness isocline, the combinations of 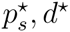 that satisfy *∂* log(*r*)_*ps*_ = 0. The red curve is the *d*^⋆^ fitness isocline. Above the black curve, selection decrease 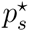. Below the black curve, selection increases 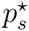. Above the red curve, selection decreases *d*^⋆^, and below it selection increases *d*^⋆^. The false positive function is set to *f*(*d*) = *ad*/(1 + *a − d*).

**Figure 2:**
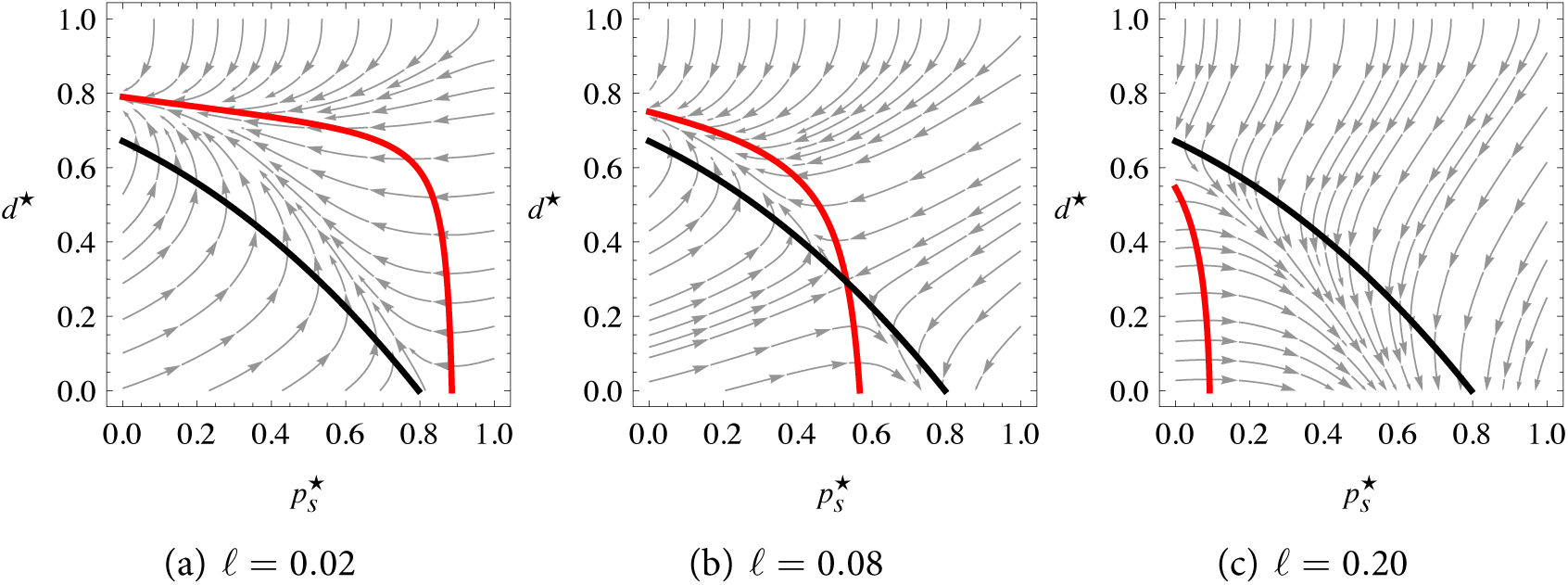
Dynamics of signal detection, as a function of the cost of detection, *ℓ*. Each figure plots the dynamics in the trait space 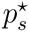 and 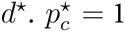 in each case, which clarifies the presentation, without loss of generality. The gray streams represent the evolutionary flow of these characters. The black curve is the combinations of 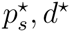 at which there is no directional change in 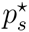. The red curve is the combinations of 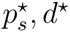 at which *d*^⋆^ does not change. The three panels vary in the direct cost of detection, *ℓ*, while holding constant *b* = 0.1, *u* = 0.3, *k* = 0.35, *w*_0_ = 1, *s* = 1, *a* = 0.1. (a) *ℓ* = 0.02: There is only one equilibrium here, where the red curve meets the left margin, at 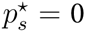. (b) *ℓ* = 0.08: There is now an unstable internal equilibrium, where the red and black curves intersect, and two stable points, at the left end of the red curve and the bottom end of the black curve. (c) *ℓ* = 0.20: The only equilibrium in this case is where the black curve meets the bottom margin, where 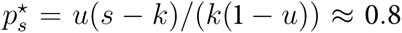 and *d*^⋆^ = 0.

Now consider each plot in Figure 2 in turn. First, when *ℓ* is very small, in panel (a), detection can both invade from zero and is stable once large. Detection invades at the point where the black curve meets the bottom axis. Since this is below the red curve in (a), selection increases detection. In this case, detection will always evolve to 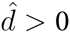 and 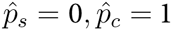. Second, when *ℓ* is intermediate, as in panel (b), detection cannot invade from zero but can be stable once large. In this case, detection may come to rest at 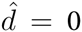 or 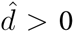, depending upon initial conditions. Third, if *ℓ* is sufficiently large, as in panel (c), detection can neither invade nor be stable. In this case, detection will always remain at 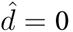.

Another way to summarize the same dynamic is to plot the best response values of *p*_*s*_, *p*_*c*_ and *d* as a function of *d*^⋆^. A best response here is the value of the trait that will maximize fitness, conditioned on the value of the other traits. Figure 3 shows these best responses. In each plot, the green, red and black curves plot the respective values of *p_s_, p_c_, d* that maximize fitness, given a population with common trait value *d*^⋆^ on the horizontal axis. I compute these by allowing *p*_*s*_ and *p*_*c*_ to go to their equilibrium values, given *d*^⋆^. This provides the values for the green (*p*_*s*_) and red (*p*_*c*_) curves. Then I compute the fitness maximizing value of *d*, conditioned on *d*^⋆^ and the best response values of *p*_*s*_ and *p*_*c*_. So to see what values of *p_s_, p_c_, d* are favored when *d*^⋆^ takes a particular value, find the value of *d*^⋆^ on the horizontal axis and then go up to find the values of *p*_*s*_ (green curve), *p*_*c*_ (red curve), and *d* (black curve) at that point.

**Figure 3:**
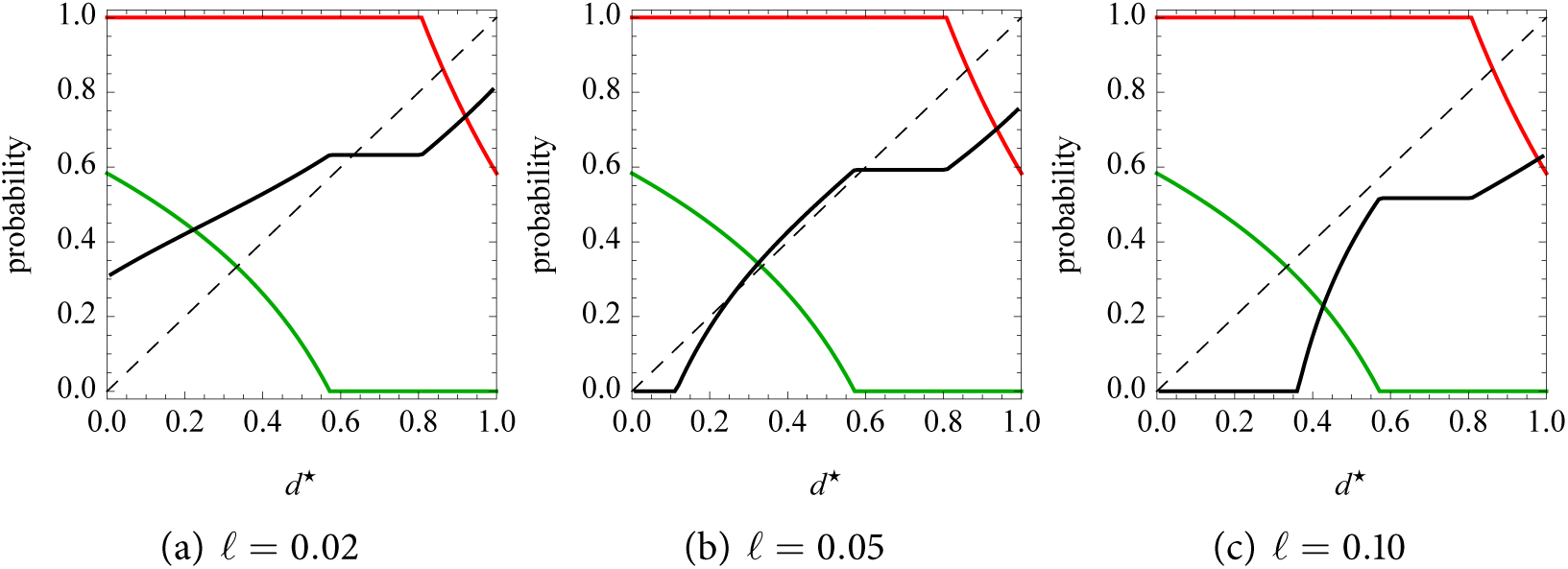
Best response values of *p_s_, p_c_, d*, as a function of *d*^⋆^ and the cost of detection, *ℓ*. In each plot above, the green, red, and black curves are the respective values of *p_s_, p_c_, d* that jointly maximize fitness, given the value of *d*^⋆^ on the horizontal axis. Where the black curve is above the diagonal, larger values of *d* increase fitness. Below the diagonal, smaller values of *d* increase fitness. Where the black curve crosses the diagonal is an equilibrium. Parameter values held at *b* = 0.1, *u* = 0.2, *k* = 0.3, *s* = 1, *w*_0_ = 1, *a* = 0.5. Plots (a), (b) and (c) vary *ℓ* so as to illustrate the same three regimes as in Figure 2. (a) Low cost and globally stable detection where the black curve crosses the diagonal. (b) Intermediate cost and a bi-stable regime. (c) High cost and globally stable *d*^⋆^ = 0.

These plots clearly illustrate a key result of the model. The stable equilibrium for 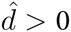, found where the black curve crosses the diagonal, always occurs where *p*_*s*_ = 0, *p*_*c*_ = 1. Behavior is perfectly separated by receiving the signal. Why should this be the case? Why can’t *d*^⋆^ stabilize where either *p*_*s*_ or *p*_*c*_ is intermediate between zero and one? The reason is complex enough to deserve it’s own section, to follow.

### 3.2. Mean fitness and the dynamics of detection

The mean fitness (log-geometric growth rate of the common type) in the population either remains constant or decreases during invasion. But near the 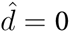 steady state, mean fitness may be both greater than or less than fitness at 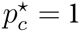. As a result, once detection evolves, the population could be either better or worse off than if no one bothered to detect environmental change. In this section, I attempt to explain these dynamics. In the process, it will become clear why *d*^⋆^ > 0 always stabilizes where 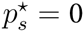 and 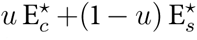.

Figure 4 illustrates the dynamics of mean fitness, as detection invades and stabilizes. Each plot in this figure has the same axes as the plots in Figure 3. In all three plots, *u* = 0.3, *k* = 0.4, and *a* = 0.1, chosen for clarity of presentation. The dashed curves represent the same kind of best response profiles as in Figure 3, with gray for *d*, light red for *p*_*c*_, and light green for *p*_*s*_. The solid curves now represent mean fitness (black), the rate of individual learning (orange), and the quality of social information (blue). The orange curves are rates of individual learning, 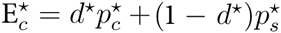, where 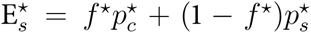 is the probability of individual learning when the environment has just changed (*t* = 0) and 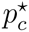 is the probability when the environment is stable (*t* > 0). The blue curves are *Q* as defined earlier. The black mean fitness curve is just the functional component of fitness, *I* + *S* − *ℓd*^⋆^, as in Equation 4. Each of these solid curves is computed for a population with *d*^⋆^ on the horizontal axis and 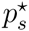 and 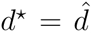 from the best response curves.

**Figure 4:**
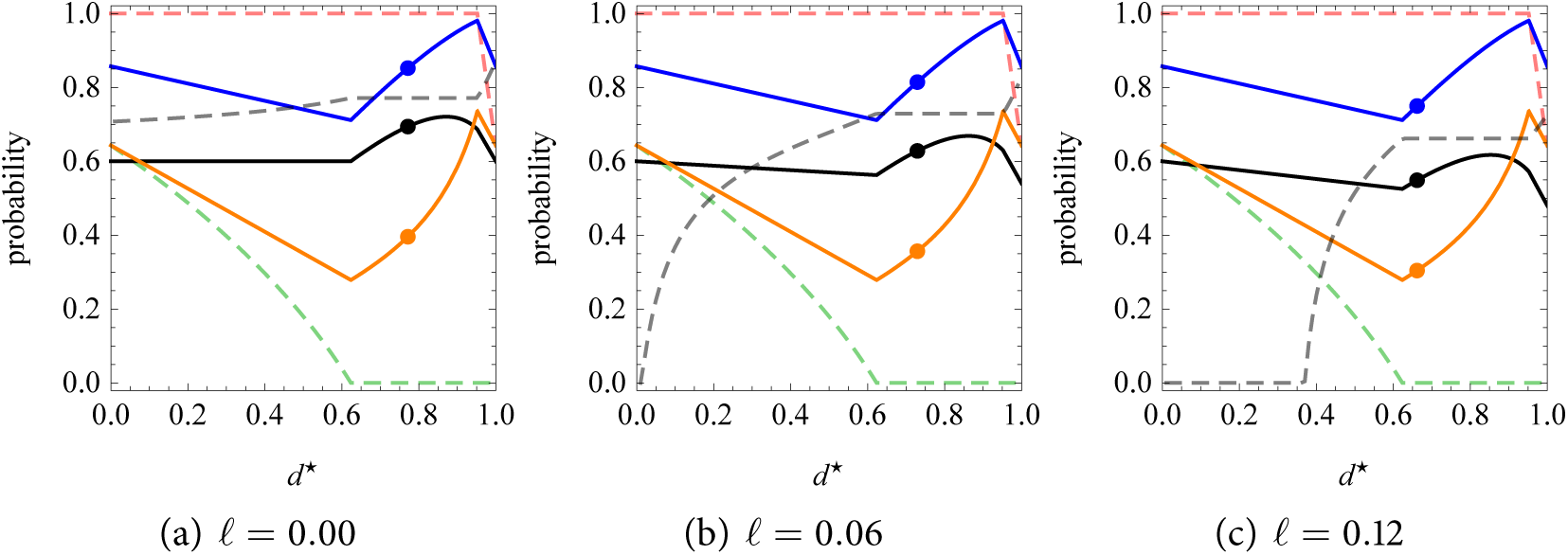
Mean fitness at the detection steady state. In each plot, the dashed gray, light red, and light green curves are the best-response *d*, *p*_*c*_, and *p*_*s*_, corresponding to the curves in Figure 3. The solid black curve is the mean fitness at *d*^⋆^ on the horizontal axis. The solid blue curve is the quality of social information, *Q*. The solid orange curve is the average rate of individual learning. The points on each curve show the values at the 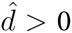 detection steady state. (a) Low cost of detection and mean fitness at 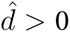 is greater than at 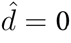. (b) Higher detection cost. Mean fitness is still higher at 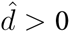, but it declines initially during invasion. (c) At very high detection cost, mean fitness at 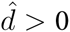 can be lower than at 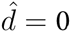. In all three plots, *u* = 0.3, *k* = 0.4, and *a* = 0.1. Dynamics are explained in the main text.

In the Supporting Information, I present mathematical analysis of rates of change of individual learning (orange) and quality of social information (blue) as they respond to changes in detection *d* and individual learning *p*_*s*_. Here, I provide a verbal summary to motivate understanding of how the average rate of individual learning and *Q* contribute to the dynamics of mean fitness.

When detection increases from zero, the mutant individual’s rate of individual learning increases. This increase results from more adaptive individual learning at *t* = 0. But it also results from more maladaptive individual learning at *t* > 0, because of false positives *f*. Selection then favors a reduction in *p*_*s*_, to both compensate for the false positives as well as reap greater *Q* resulting from the spread of detection in the population. But this reduction in *p*_*s*_, once it spreads through the population, reduces both the overall rate of individual learning (orange) as well as the quality of social information *Q* (blue) in the population. This reduction in the quality of social information cancels any mean fitness benefit of detection. So during this phase of the dynamics, the rate of individual learning in the population declines, but accompanying decline in the quality of social information means the population experiences no average fitness benefit from avoiding individual learning. This dynamic is very much like the one that generates constant mean fitness in many models of this kind (Boyd and Richerson 1995).

However, mean fitness is not always constant during this phase of the dynamics. If detection is individually costly (*ℓ* > 0), as in the middle plot in Figure 4, mean fitness (black curve) will actually decline as detection invades. This decline does not prevent detection from invading, however, because the mutant is playing the market and does experience a relative fitness benefit initially. It is only once the rest of the population catches up that the quality of social information declines and reduces mean fitness. Thus in such scenarios, detection invades but actually makes the population worse off.

Once detection is high enough, however, 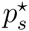 reaches zero and cannot be reduced any further. Any further increases in detection now increase the rate of individual learning, because false positives at *t* > 0 cannot be compensated for by reducing *p*_*s*_. The orange curves rise. As a result, the quality of social information also rises, producing a population benefit of more accurate behavior to imitate. Detection *d*^⋆^ can continue to increase, raising both the quality of social information and mean fitness, until the rate of false positives (*f*^⋆^) satisfies Equation 8. At that point, selection stabilizes 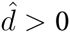.

Mean fitness at 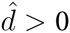 can therefore be higher than that at 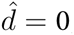, as in the lefthand and middle plots in Figure 4. But if the costs of detection *ℓ* are large enough, as in the righthand plot, then mean fitness at 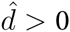 may actually be lower than that at 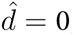. The same dynamic as above is at work here, but now the direct cost of detection is large enough that the increase in mean fitness near 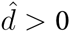 cannot overcome it.

This dynamic helps to understand why 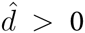 can only stabilize where 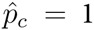 and 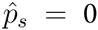 and why mean fitness can only increase in the same region. Until 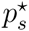 reaches its minimum, selection can compensate for an increase in false positives by both reducing individual learning when *t* > 0. This compensation has the consequence of spoiling the quality of social information, erasing fitness gains of invaders. This is similar to the dynamic in Rogers’ model and many similar models (Rogers 1988, Boyd and Richerson 1995), in which invading social learners eventually spoil the quality of social information, erasing any fitness gains for the population. However in this model, once 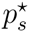 cannot be further reduced to compensate for increasing *f*, invaders improve the quality of social information. Now mean fitness (black curves in Figure 4) will rise, both because of (1) the direct benefits of detection allowing individuals to allocate expensive individual learning to when it is needed most and (2) the population side effect of improving the quality of social information.

Further increase in *d*^⋆^ beyond 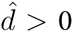 would increase mean fitness, as can be seen by the location of the black points in Figure 4 always lying to the left of the peak of the black curve. However, in the absence of some factor like kin selection (in a non-viscous population structure), natural selection will not maximize mean fitness in this model. Detection is individually costly, but produces a population benefit by increasing the quality of social information, near steady state. This is a kind of collective action dilemma, similar to the basic collective action dilemma embodied in individual and social learning: individual learning is individually costly, but produces population benefits. Ironically, kin selection would increase detection at 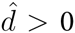, but also narrow the conditions that allow detection to invade, because now the depression of mean fitness during invasion (*ℓ* > 0, as in the middle plot) would reduce inclusive fitness.

## 4. Discussion

### 4.1. When does detection evolve?

Detection of recent environmental change evolves under many circumstances that favor reliance on social learning at all. Selection further favors detection when individual learning is costly relative to the rate of environmental change, detection is efficient (not too many false positives), and detection is not too costly. These conclusions can be read directly from Condition 7. To understand them, consider that the initial problem detection solves is allocation of costly individual learning to generations immediately following environmental change. Detecting change allows individuals to likewise allocate more social learning to generations in which the environment is stable. But if individual learning is cheap relative to the rate of environmental turnover, then little social learning is favored even when the environment is stable. At the extreme where the rate of change *u* is larger than the effective costs of individual learning *k/s*, detection can never invade the population.

However, detection does invade over a broad range of parameter values. This is most evident perhaps in the sensitivity plots in the Supporting Information (Figures 5 and 6). The intuition behind this result is that, whenever substantial social learning is favored in the absence of detection, as it routinely is in such models, there will be a basic allocation problem that can be addressed by detecting recent environmental change. Social learning is a risky, high variance learning strategy, relative to individual learning. Just after a change in the environment, all social learning results in zero probability of acquiring adaptive behavior. This effect is very stark in this kind of model, because all adaptive information is lost when the environment changes. But the general principle appears robust, as it remains even in cumulative culture models in which some adaptive behavior can persist (McElreath 2010, e.g.). Detection reduces the fitness variance of social learning, by allocating more of it to when it is safest to use.

**Figure 5.**
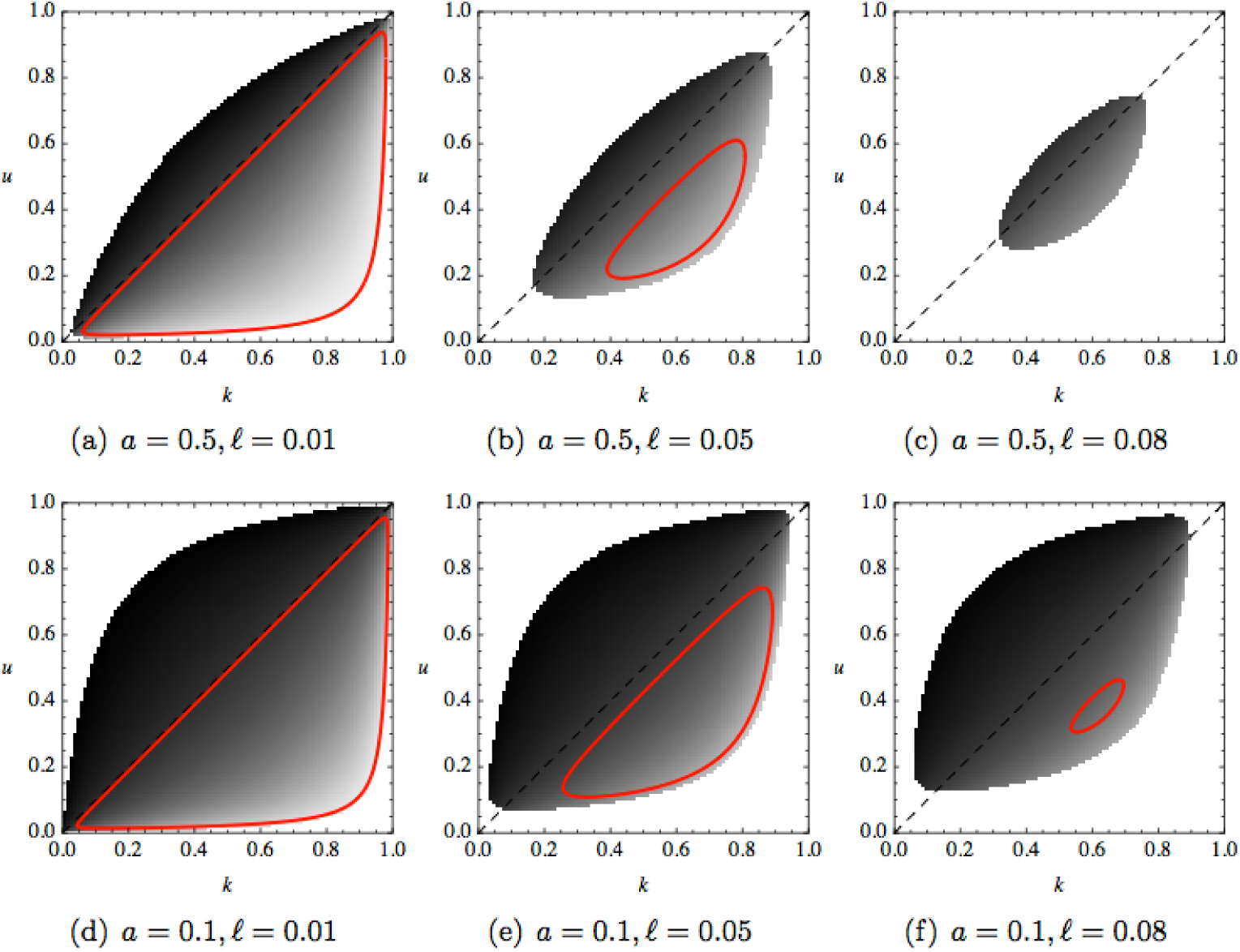
Sensitivity of invasion and stability to the parameters *k*, the cost of individual learning, and *u*, the probability of environmental change. In each plot, the red boundary contains the combinations of *k, u* that allow detection to invade. The shaded region shows all combinations of *k, u* at which detection is stable when common. Darker shading indicates higher equilibrium detection, 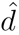, with pure black representing 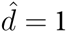 and pure white 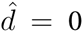. Inside the shaded region, 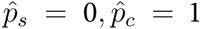. Outside the shaded region, 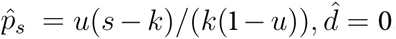. Note that detection can be stable at high values even when there are no combinations of *k, u* that allow invasion, as in panel (c).

**Figure 6.**
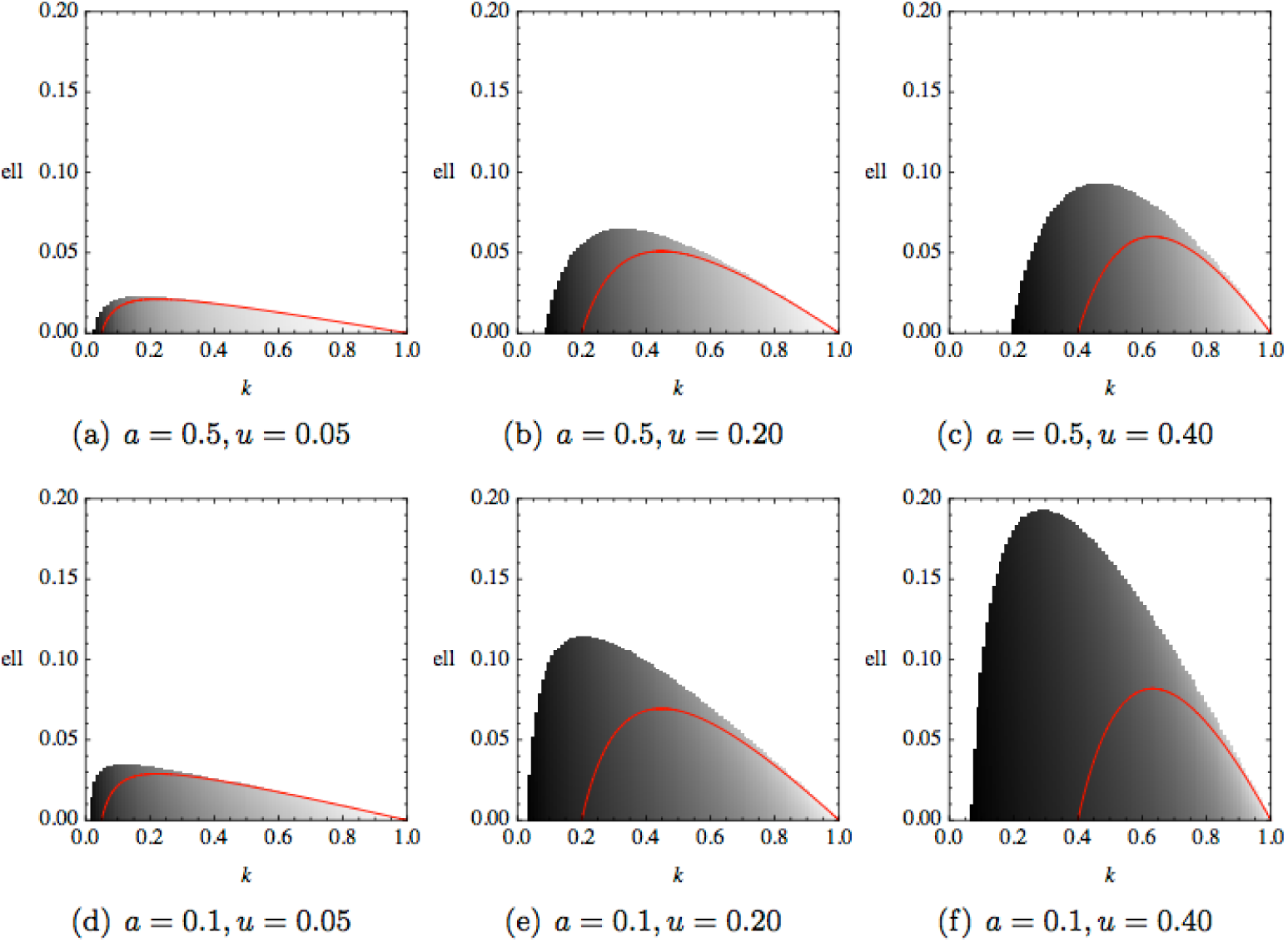
Sensitivity of invasion and stability to the parameters *k*, the cost of individual learning, and *ℓ*, the direct cost of detecting environmental change. In each plot, the red boundary contains the combinations of *k, ℓ* that allow detection to invade. The shaded region shows all combinations of *k, ℓ* at which detection is stable when common. Darker shading indicates higher equilibrium detection, 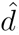, with pure black representing 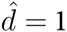 and pure white 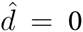. Note the scale of the vertical axis, which unlike the horizontal, only extends to 0.20.

### 4.2. How does detection work?

Of course these results must overstate the probability of detection’s evolution, because the model assumes a cue of environmental change is available and that the organism can discover it. The nature of such cues is left abstract in the model, but the sensory abilities of the organism and structure of the population must constrain the possibilities. Potentially general cues of recent environmental change may include poor health or fertility of conspecifics. Organisms with sophisticated communication, like humans, may also detect change by paying attention to reflections of older individuals. For example, contemporary arctic peoples possess elaborate, ecologically-accurate models of their environments. When events fall outside past patterns, younger individuals can and do benefit from listening to such observations (Fox 2002, Weatherhead *et al.* 2010). Under this view of detection, the costs are in attention and processing, being possibly quite small.

Another idea is that naive individuals learn socially first and that the cost of detection in this model represents the cost of trying out socially-learned behavior. This would make detection here like “critical social learning” (Boyd and Richerson 1996, Enquist *et al.* 2007), in which individuals first learn socially, have a chance of diagnosing maladapted behavior, and can finally use individual learning as a last resort. Like detection in this model, critical social learning can be stable even when it cannot invade, and it can raise mean fitness. Unlike detection however, critical social learning models currently contain no false positives—although there is transmission error, critical social learners never mistake adaptive behavior for maladaptive. The detection model here in contrast contains inherent risk of excessive individual learning, because of mistaken diagnosis. But the key point is that critical social learning may be an alternative mechanism for detecting temporal environmental change, one that is rather accessible to evolution. Its dynamics will be different from the use of cues specifically tied to ecological change, such as those mentioned in the previous paragraph, but may nevertheless coevolve.

### 4.3. What are detection’s effects?

Once detection does invade, the distribution of individual and social learning through time changes. A dominant result is that a population that invests in detecting recent change will exhibit less individual learning overall. It will also exhibit much less individual learning during periods of environmental stability.

This allocation of individual learning to periods just following a change results in a rapid increase in the frequency of adaptive behavior, just after a change in environment. But it also results in a very slow increase afterwards. As a result, the frequency of adaptive behavior in stable environments may not look very different, after detection evolves. However, the frequency of adaptive behavior recently following a change in the environment will look quite different, showing a rapid increase. In the end, a snapshot of a population in which detection has evolved will show a higher reliance on social learning compared to what evolves in the absence of detection.

All of these population dynamics combine to allow average fitness, or the expected population growth rate, to rise after detection evolves. This increase in mean fitness is usually quite modest in this model, much smaller than that demonstrated from cumulative culture models. However the increase appears for much the same reason: detection allows individual and social learning to work together synergistically, rather than competitively (Boyd and Richerson 1995). Learning individually just after a change in the environment both helps those doing the learning as well as the social learners in following time periods. However, the benefit to subsequent generations does not lead to natural selection reducing individual learning, because the benefit was produced at a time that social learning produces no fitness increase at all. Therefore the “freeloading” by social learners during stable periods does not threaten to erode the public good provided by individual learners just after a change. But if detection is very costly or individual learning is sufficiently cheap, then the change in mean fitness may be very small or even negative.

### 4.4. Model variations

I have chosen this model’s features because they represent a central case for analysis, one comparable to existing theory. However, all models are necessarily special, and so future work should address other model assumptions.

#### Spatial environmental variation

It has long been recognized (Hedrick *et al.* 1976) that spatial and temporal environmental variation produce different selection regimes. This is just as true for gene-culture coevolutionary models (McElreath *et al.* in press, Nakahashi *et al.* in press). “Detection” in a spatial variation context would mean the population is sub-divided into a number of patches. A different behavior is optimal in each patch. Individuals can evolve different learning strategies depending upon whether or not they are recent immigrants to a local patch. A first conjecture is that selection will favor greater reliance on social learning for recent immigrants. It would also be possible to examine whether selection may favor residents’ ignoring immigrants, when choosing models to learn from. An important question to ask of such a model is whether adjusting use of social learning depending upon migration status will allow unbiased social learning to maintain cultural variation, even when it cannot in traditional models.

Spatial and temporal variation may also interact in unanticipated ways. Such interactions have been well-explored in the study of dispersal (Schreiber 2010, for a recent example), but the importance of these phenomena is potentially much more general (Williams and Hastings 2011).

#### Other learning strategies

Unbiased social learning, in which a single target of learning is chosen independent of its behavior, is a very special case. The most-discussed alternatives includes conformist transmission (Boyd and Richerson 1985, Henrich and Boyd 1998) and some kind of payoff or success or prestige biased transmission (Boyd and Richerson 1985, Henrich 2001, McElreath *et al.* 2008).

In the case of conformist transmission, recent debates over whether or not selection will favor it provide a natural opening to consider contingent use. Conformist transmission was original studied as an adaptation to spatial environmental variation (Boyd and Richerson 1985, Henrich and Boyd 1998). Wakano and Aoki (2007) later studied a model of conformist learning in which there was only temporal variation, finding that the conditions that favored conformist transmission were very restrictive. Some of the same authors have more recently studied conformist transmission under both temporal and spatial variation, confirming the original intuition that it is an adaptation to spatial variation (Nakahashi *et al.* in press). McElreath *et al.* (in press) have recently shown that a mix of temporal and spatial variation can also favor a strong reliance on conformist transmission, as in Henrich and Boyd’s model. Finally, a recent Bayesian model (Perreault *et al.* in press) demonstrates a robust reliance on conformist transmission, even when the environment varies only temporally.

An explicit consideration of contingent use of conformist transmission as a function of cues of environmental change and migration status should help to unify this literature. It would also help in interpretation of experimental results. All of the existing experimental and quasi-experimental studies of social learning contained analogues of only temporal environmental variation. While conformist transmission has been found in some of these cases (Kameda and Nakanishi 2002, McElreath *et al.* 2005, 2008), it has not always been found (Eriksson *et al.* 2007, Eriksson and Coultas 2009). Experiments that allow for the analogue of spatial variation should provide cleaner tests.

#### Learning costs

When there are multiple domains of behavior to be learned, and the costs of learning vary among them, how will selection design learning? Since the problems the organism needs to solve may change across space and time, it is problematic to assume that there can be a genetic locus controlling reliance on social learning in each domain. Should an organism attempt to estimate a cost of individual learning in each domain, or rather adapt to a fitness-weighted average of the domains?

## 5. Conclusion

To return to the problem that motivated this model: How do these results reflect on the interpretation of social learning experiments? If human or other animal participants do have strategies attuned to cues of environmental change, we will need to consider whether or not our experiments accidentally include too many or too few such cues. For example, in the typical experiment, all participants all equally naive at the start. This may function as a social cue of recent environmental change, in that it favors increased reliance on individual learning. On the other hand, experiments that provide participants with no way to detect changes in the underlying payoffs may accidentally provide cues of environmental stability. In the end, explicitly designing both laboratory and field studies with contingent strategy use in mind will provide clearer tests of theory.

### supporting Information

#### Derivation of *q*_*t*_

First, note that just after a change in the environment, *q*_*t*_ resets to *q*_0_ = 0. One generation after a change in the environment, *t* = 1, the expected chance of acquiring adaptive behavior via social learning is:

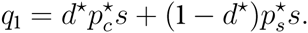

Here’s how to motivate the expression above. The only way to acquire adaptive behavior this soon after a change in the environment is to target an individual who learned individually in the last generation. A proportion 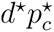 of the previous generation correctly noticed the change in the environment and chose to learn individually with chance 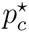. The remaining proportion 1 − *d*^⋆^ failed to detect the change in the environment and continued to learn individually with chance 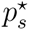.

Next, in any generation *t >* 1, the quality of social information is given by:

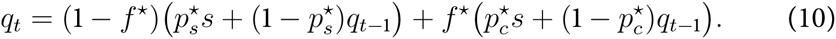

Here’s how to motivate this expression. A proportion 1 − *f*^⋆^ of the population did not wrongly conclude that the environment changed recently. They learn individually 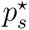 of the time and update socially 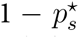 of the time. Social updating leads to successful acquisition of adaptive behavior with probability *q*_*t*−1_. The rest of the population, a proportion *f*^⋆^, thinks the environment just changed and updates accordingly.

The above recurrence equation for *q*_*t*_ (Equation 10) can be solved explicitly for a function *q*_*t*_ that is not a function of *q*_*t*−1_. It is a linear recurrence and so several methods exist. I used the *Ansatz* method of guessing the form and proving it was correct. The resulting function is:

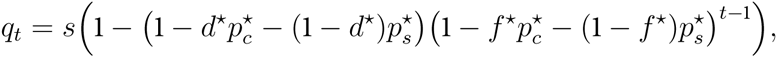

for all *t* ≥ 1.

#### Invasion and stability conditions for any function *f*(*d*)

To find the conditions for this equilibrium to exist and be stable, we can observe that the dynamics of 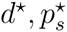 are governed by two null clines, where the change in each state variable is zero, as a function of 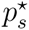 (Figure 2). Both null clines are convergent, in the sense that dynamics take each state variable closer to its null cline. Thus if *d*^⋆^ is plotted on the vertical axis and 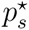 on the horizontal axis, then the system moves up when it is below the null cline for *d*^⋆^ and left when it is to the right of the null cline for 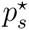. Because these null clines cross only once, we can inspect the four end points on both the left and bottom axes to summarizes the dynamics of the system.

Consider first the bottom axis, where *d*^⋆^ = 0. The 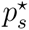 null cline will always intersect the bottom axis at 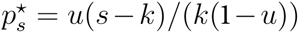. Then depending upon whether the *d*^⋆^ null cline lies left or right of this point determines whether detection can invade. The null cline for *d*^⋆^ must lie to the right of the null cline for 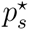, for *d*^⋆^ to increase. Otherwise the system at the invasion point 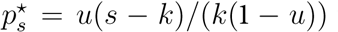 will be above the *d*^⋆^ null cline and decrease. If instead the *d*^⋆^ null cline is right of 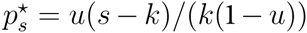, the system will lie below the *d*^⋆^ null cline and therefore *d*^⋆^ will increase. So the condition for *d*^⋆^ to increase from the point 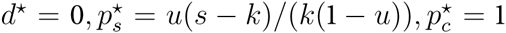 is given by asking when the value of 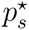 that makes *∂*log(*r*)_*d|d*^⋆^=0_ = 0 is greater than 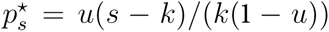. When instead this point is lower than 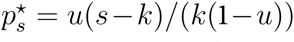, detection cannot invade from zero. Reducing this condition tells us that stability at *d*^⋆^ = 0 requires either that *k* ≤ *su* or *s* ≤ *k*, or when *k* > *su* and *s* > *k*, it requires:

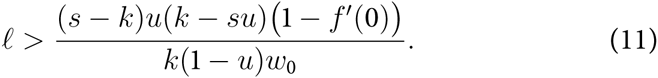

When this condition is satisfied, detection cannot increase from zero.

The second condition for the internal unstable equilibrium to exist is that the null cline for *d*^⋆^, along the left axis where 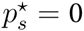, be greater than the null cline for 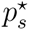 (Figure 2). This reduces to the condition:

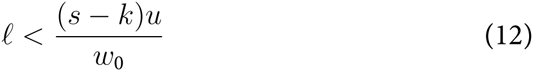

and

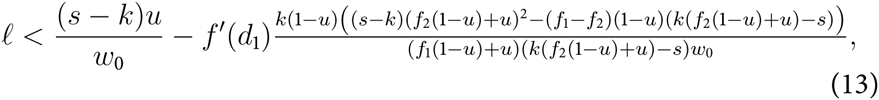

where *d*_1_ is the value of *d*^⋆^ that satisfies 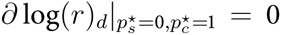 and *d*_2_ the value of *d*^⋆^ that satisfies 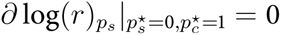. The symbol *f*_1_ ≡ *f* (*d*_1_) and *f*_2_ ≡ *f*(*d*_2_). For condition 13 to be less than condition 12, it is also necessary that:

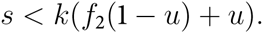

In summary, when detection cannot invade (condition 11 is satisfied) but is stable when large (condition 13 is satisfied), the dynamics contain an internal unstable equilibrium. This proves that the signal detection equilibrium with stable *d*^⋆^ > 0 and 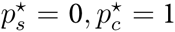 is stable for a broader range of values than will allow *d*^⋆^ > 0 to invade from *d*^⋆^ = 0.

The dynamics of the system can also be summarized in terms of these two conditions. There are three possible combinations. First, detection can invade when rare and be stable when common. This requires that condition 11 be false and condition 13 true. Second, detection cannot invade when rare but can be stable once at a large enough value. This holds when condition 11 is true and condition 13 is true. Third, detection can neither invade nor be stable once large. This holds when condition 11 is true and condition 13 is false.

#### Sensitivity plots

It is much easier to appreciate the effects of the parameters on invasion, stability, and equilibrium detection rate by using sensitivity plots. In Figures 5 and 6, I illustrate how the parameters influence the relative sizes of the invasion and stability conditions, as well as the steady state detection rate. These plots all use the box hyperbola ROC function, *f*(*d*) = *ad*/(1 + *a − d*).

Figure 5 plots evolutionary outcomes of the model, within the parameter space defined by *k*, the cost of individual learning, and *u*, the instability of the environment. Each of the six plots varies the accuracy of detection, *a*, and the direct cost of detection, *ℓ*. All other parameters are held constant at *w*_0_ = 1, *s* = 1. The red regions enclose all combinations of *k, u* that lead *d*^⋆^ to increase from zero. These are the invasion regions. The shaded regions enclose all combinations of *k, u* for which detection can be stable once large enough. These are the stability regions. The degree of shading in the stability regions represents the amount of detection at equilibrium, for each combination of *k, u*. Pure black represents 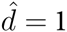 while pure white represents 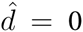.

In every case, the red invasion region does not extend above the diagonal where *k/s* = *u*. When *u* > *k/s* and *d*^⋆^ = 0, individuals are already using individual learning 100% of the time. If individuals had a flawless signal of environmental change, then selection would favor detection and using social learning when the environment is stable. But the signal is never perfect. Instead, attempts to detect stability always lead to some erroneous decisions to learn socially. When *u* is large, the probability the environment has not changed will be small and comparable to the rate of mistakes in concluding that the environment has not changed. Since individual learning is so cheap, when *u > k*, the risks do not outweigh the costs, and detection can never invade.

For combinations of small *u* and large *k* (lower-right corner of each plot), invasion is similarly impossible. In these regions, very little individual learning is favored, because of its high cost and the infrequency of change in the environment. For example, where *u* = 0.1 and *k* = 0.9, the stable proportion of individual learning is 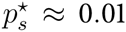. Now the environment doesn’t change much, and attempts to detect change will generate errors at a rate comparable to the probability of true change.

Detection can be stable over a larger region, however, above *u* = *k/s*. Even though detection cannot invade where *u > k/s*, it may be stable once large. Once detection is accurate enough, it allows individuals to allocate more individual learning to when it is most needed and simultaneously reduce their overall reliance on individual learning, which is expensive.

A similar phenomenon does not appear in the corner where *k* is large and *u* is small, because so little individual learning is favored there that detection cannot reduce fitness costs much by reducing overall reliance on individual learning—there just isn’t much reliance to reduce. As a result, the stability regions can extend far above the *k/s* = *u* invasion boundary, but not very far beyond the lower-right of it.

In the top row of Figure 5, accuracy of detection is set fairly low, to *a* = 0.5. At this accuracy, in order to correctly detect 50% of all environmental changes, an individual would suffer false alarms 25% of the time. In order to correctly detect 90% of changes, an individual would suffer false alarms 75% of the time! The three plots in this row vary the direct cost of detection, *ℓ*. Both the invasion and stability regions shrink rapidly with increases in the direct cost of detection.

In the bottom row of Figure 5, accuracy of detection is set fairly high, to *a* = 0.1. And at this accuracy, 50% true detection implies a false alarm rate of only 8.3%. A 90% detection rate implies a 45% false alarm rate. When accuracy is this high, changes in the direct cost of detection, *ℓ*, have much less of an effect on the stability region.

It is easier to appreciate the effect of *ℓ* by holding *u* constant and varying *ℓ* and *k*. I do this in Figure 6, for three values of *u* (0.05,0.20,0.40) and the same two values of *a* as in Figure 5 (0.5,0.1). In the space defined by *ℓ* and *k*, the red invasion region rises above zero where *k* = *u*. This is the boundary on the diagonal in Figure 5. We can now see, however, that the invasion region extends all the way to the right, provided that *ℓ* = 0. For *ℓ >* 0, intermediate values of *k* have the largest invasion potential. This reflects the same tradeoffs that I explain for the *u, k* parameter space.

The effect of the accuracy of individual learning, *s*, is to compress the space defined by *k*. The true dimension of the cost of individual learning is *k/s*, not *k*. When *s* = 1, as in the previous figures, *k* summarizes the cost of individual learning. But for smaller *s*, the horizontal axis is effectively compressed, otherwise leaving the geometry unchanged.

#### Dynamics of *L* and *Q*

The expected rate of individual learning, as a function of *d, p*_*c*_, *p*_*s*_ is:

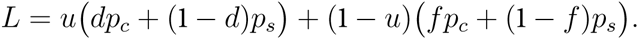

This expression is for a mutant, but since individual learning is asocial, the rate for the population is analogous, using 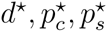

We want to prove that *L* increases with *d*, in order to demonstrate that increasing detection increases individual learning on average. The rate of change in *L* as a function of *d* is, via the chain rule:

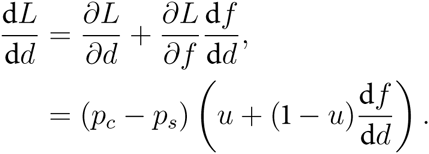

Since d*f*/d*d >* 0 and *p_c_ − p_s_ >* 0, as conditions for *d >* 0 to invade in the first place, d*L*/d*d >* 0.

Now consider the rate of change in *L* as a function of *p*_*s*_. This is:

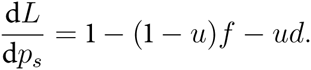

And this is also always positive, for any 0 < *u* < 1 and 0 < *f* < *d* < 1. Increasing *p*_*s*_ increases *L*, and so decreasing *p*_*s*_ decreases *L*, explaining the negative trend for *L* as *p*_*s*_ approaches zero.

Now consider the change in the quality of social information, *Q*. Again via the chain rule, the rate of change in *Q* as a function of *d*^⋆^ is:

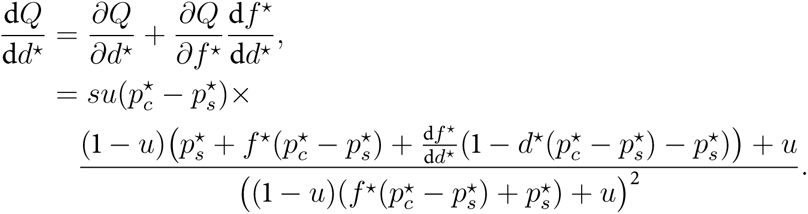

While this appears complicated, it is always positive for any 0 < *f*^⋆^ < *d*^⋆^ < 1 and 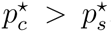. It’s worth noting also that the above is proportional to *u*, because detection improves *Q* partly by focusing more individual learning on time periods where *t* = 0. This increases *Q* for all *t >* 0, as a consequence. The more common change in the environment, the more detection helps *Q*.

*Q* also increases with 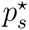. The rate of change is:

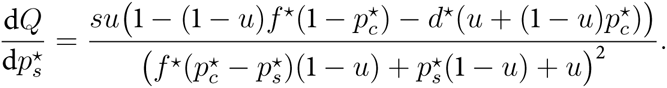

And this is also positive, for all 0 < *f*^⋆^ < *d*^⋆^ < 1. Therefore as 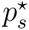 decreases, *Q* decreases.

In this model, the evolution of detection of environmental change can both increase or decrease mean fitness. The analytical conditions for these outcomes are complex functions of every variable in the model. However, considering the special case where *ℓ* = 0 does provide some qualitative insight. The condition for expected fitness at the 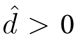 steady state to exceed fitness at 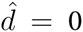 is:

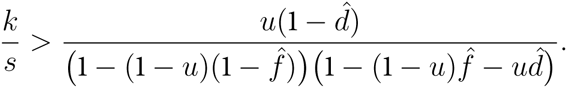

As the costs of individual learning *k* increase, mean fitness at 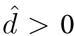 increases. This results from the population being able to save costs of expensive learning, while still producing quality social information, by allocating necessary individual learning to when it is really needed, when *t* = 0. A major opposing force is the rate of change *u*, which reduces mean fitness at 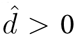. As *u* increases, the population spends less and less time at *t >* 0, and so reaps less benefit from any improvements in social information *Q*. Finally, the slower *f* increases with *d*, the higher mean fitness at 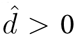

## References

Altenberg, L. 2012. Resolvent positive linear opperator exhibit the reduction phenomenon. Proceedings of the National Academy of Sciences USA 109:3705–3710.

Altenberg, L. in press. The evolution of dispersal in random environments and the principle of partial control. Ecological Monographs.

Baum, W. M., P. J. Richerson, C. M. Efferson, and B. M. Paciotti. 2004. Cultural evolution in laboratory microsocieties including traditions of rule giving and rule following. Evolution and Human Behavior 25:305–326.

Boyd, R., and P. J. Richerson. 1995. Why does culture increase human adaptability? Ethology and Sociobiology 16:125–143.

Boyd, R., and P. Richerson. 1996. Why culture is common, but cultural evolution is rare. Proceedings of the British Academy 88:77—93.

Boyd, R., and P. J. Richerson. 1985. Culture and the Evolutionary Process. Chicago: Univ Chicago Press.

Cavalli-Sforza, L. L., and M. W. Feldman. 1981. Cultural transmission and evolution: a quantitative approach. Princeton: Princeton University Press.

Cohen, D. 1966. Optimizing reproduction in a randomly varying environment. Journal of Theoretical Biology 12:119–129.

Cohen, D. 1967. Optimizing reproduction in a randomly varying environment when a correlation may exist between the conditions at the time a choice has to be made and the subsequent outcome. Journal of Theoretical Biology 16:1–14.

DeWitt, T. J., A. Sih, and D. S. Wilson. 1998. Costs and limits of phenotypic plasticity. Trends in Evolution and Ecology 13:77–81.

Efferson, C., P. J. Richerson, R. McElreath, M. Lubell, E. Edsten, T. M. Waring, B. Paciotti, and W. Baum. 2007. Learning, Productivity, Noise: An Experimental Study of Cultural Transmission on the Bolivian Altiplano. Evolution and Human Behavior 28:11–17.

Efferson, C., R. Lalive, P. J. Richerson, R. McElreath, and M. Lubell. 2008. Conformists and mavericks: The empirics of frequency-dependent cultural transmission. Evolution and Human Behavior 29:56–64.

Enquist, M., K. Eriksson, and S. Ghirlanda. 2007. Critical Social Learning: A Solution to Rogers’s Paradox of Nonadaptive Culture. American Anthropologist 109:727–734.

Eriksson, K., and J. C. Coultas. 2009. Are people really conformist-biased? An empirical test and a new mathematical model. Journal of Evolutionary Psychology 7:5–21.

Eriksson, K., M. Enquist, and S. Ghirlanda. 2007. Critical points in current theory of conformist social learning. Journal of Evolutionary Psychology 5:67–87.

Eriksson, K., and P. Strimling. 2009. Biases for acquiring information individually rather than socially. Journal of Evolutionary Psychology 7:309–329.

Fox, S. 2002. “These are things that are really happening”: Inuit perspectives on the evidence and impacts of climate change in Nunavut. Pages 12–53 of: Krupnik, I., and D. Jolly (eds), The Earth is Faster Now: Indigenous Observations of Arctic Environmental Change. Fairbanks, Alaska: Arctic Research Consortium of the United States.

Getty, T. 1996. The maintenance of phenotypic plasticity as a signal detection problem. The American Naturalist 148:378–385.

Green, D. M., and J. A. Swets. 1966. Signal detection theory and psychophysics. New York, NY: John Wiley and Sons Inc.

Hedrick, P. W., M. E. Ginevan, and E. P. Ewing. 1976. Genetic polymorphism in heterogenous environments. Annual Review Ecology Systematics 7:1–32.

Henrich, J. 2001. Cultural Transmission and the Diffusion of Innovations: Adoption dynamics indicate that biased cultural transmission is the predominate force in behavioral change and much of sociocultural evolution. American Anthropologist 103:992–1013.

Henrich, J., and R. Boyd. 1998. The evolution of conformist transmission and between-group differences. Evolution and Human Behavior 19:215–24.

Henrich, J., S. Heine, and A. Norenzayan. 2010. The Weirdest People in the World? Behavioral and Brain Sciences.

Jacobs, R. C., and D. T. Campbell. 1961. The perpetuation of an arbitrary tradition through several generations of a laboratory microculture. The Journal of Abnormal and Social Psychology 62:649–658.

Kameda, T., and D. Nakanishi. 2002. Cost-benefit analysis of social/cultural learning in a non-stationary uncertain environment: An evolutionary simulation and an experiment with human subjects. Evolution and Human Behavior 23:373–393.

Levins, R. 1968. Evolution in Changing Environments. Princeton: Princeton University Press.

McElreath, R. 2004. Social learning and the maintenance of cultural variation: An evolutionary model and data from East Africa. American Anthropologist 106:308–321.

McElreath, R. 2010. The coevolution of genes, innovation and culture in human evolution. Pages 451–474 of: Silk, J., and P. Kappeler (eds), Mind The Gap: Tracing the Origins of Human Universals. Springer.

McElreath, R., and P. Strimling. 2008. When natural selection favors learning from parents. Current Anthropology 49:307–316.

McElreath, R., M. Lubell, P. J. Richerson, T. Waring, W. Baum, E. Edsten, C. Efferson, and B. Paciotti. 2005. Applying evolutionary models to the laboratory study of social learning. Evolution and Human Behavior 26:483–508.

McElreath, R., A. V. Bell, C. Efferson, M. Lubell, P. J. Richerson, and T. Waring. 2008. Beyond existence and aiming outside the laboratory: Estimating frequency-dependent and payoff-biased social learning strategies. Philosophical Transactions of the Royal Society B 363:3515–3528.

McElreath, R., A. Wallin, and B. Fasolo. in press. The Evolutionary Rationality of Social Learning. In: Hertwig, R., U. Hoffrage, and the ABC Research Group (eds), Simple Heuristics in a Social World. Oxford University Press.

McNamara, J. M., and S. R. X. Dall. 2011. The evolution of unconditional strategies via the ‘multiplier effect’. Ecology Letters 14:237–243.

Morgan, T. J. H., L. E. Rendell, M. Ehn, W. Hoppitt, and K. N. Laland. 2012. The evolutionary basis of human social learning. Proceedings of the Royal Society London B 279:653–662.

Nakahashi, W., J. Wakano, and J. Henrich. in press. Adaptive social learning strategies in temporally and spatially varying environments. Human Nature.

Perreault, C., C. Moya, and R. Boyd. in press. A Bayesian approach to the evolution of social learning. Evolution and Human Behavior.

Rendell, L., R. Boyd, D. Cownden, M. Enquist, K. Eriksson, M. W. Feldman, L. Fogarty, S. Ghirlanda, T. Lillicrap, and K. N. Laland. 2010. Why copy others? Insights from the social learning strategies tournament. Science 328:208–213.

Rogers, A. R. 1988. Does biology constrain culture? American Anthropologist 90:819–831.

Schreiber, S. J. 2010. Interactive effects of temporal correlations, spatial heterogeneity and dispersal on population persistence. Proceedings of the Royal Society London B 277:1907–1914.

Sih, A., M. C. O. Ferrari, and D. J. Harris. 2011. Evolution and behavioural responses to human-induced rapid environmental change. Evolutionary Applications 4:367–387.

Tufto, J. 2000. The evolution of plasticity and nonplastic spatial and temporal adaptations in the preseence of imperfect environmental cues. The American Naturalist 156:121–130.

Wakano, J. Y., and K. Aoki. 2007. Do social learning and conformist bias coevolve? Henrich and Boyd revisited. Theoretical Population Biology 72:504–512.

Weatherhead, E., S. Gearheard, and R. G. Barry. 2010. Changes in weather persistence: Insight from Inuit knowledge. Global Environmental Change 3:523–528.

Williams, P. D., and A. Hastings. 2011. Paradoxical persistence through mixed-system dynamics: towards a unified perspective of reversal behaviours in evolutionary ecology. Proceedings of the Royal Society London B 278:1281–1290.

